# 2-monopalmitin, but not 1-monopalmitin, enhances hypothalamic leptin responsiveness, energy balance, and glucose homeostasis under overnutrition

**DOI:** 10.1101/2024.08.18.608432

**Authors:** Nozomi Takahashi, Mutsumi Ikeda, Yukiko Yamazaki, Yui Funatsu, Tamae Shiino, Aoi Hosokawa, Kentaro Kaneko

**Author notes:** Address correspondence to: Kentaro Kaneko, Department of Agricultural Chemistry, School of Agriculture, Meiji University, 1-1-1, Higashimita, Tama-ku, Kawasaki-shi, Kanagawa, 214-8571, Japan. These authors contributed equally to this work (Nozomi Takahashi, Mutsumi Ikeda).

## Abstract

Nutrient excess, a major driver of obesity, diminishes hypothalamic responses to exogenously administered leptin, a critical hormone for energy balance. Here, we found that 2-monopalmitin, but not 1-monopalmitin or palmitic acid, enhances hypothalamic leptin responsiveness in *ex vivo* brain slices. Centrally administered 2-monopalmitin markedly restored the leptin-induced suppression of food intake and reduction of body weight in diet-induced obese mice. Peripherally administered 2-monopalmitin also enhanced the anorectic effect of centrally administered leptin. Furthermore, daily 2-monopalmitin treatment protected against diet-induced body weight gain, and the energy expenditure of 2-monopalmitin-treated mice was significantly enhanced in a leptin-dependent manner. We also demonstrated that 2-monopalmitin lowered blood glucose levels, improved glucose and insulin tolerance, and protected mice against HFD-induced peripheral insulin resistance at the cellular and whole-body levels. Finally, treatment with 2-monopalmitin protected against LPS-induced leptin resistance, and decreased the hypothalamic levels of SOCS3, an inhibitor of leptin actions, and inflammatory cytokines. Altogether, our results showed that 2-monopalmitin in the brain, but not 1-monopalmitin or palmitic acid, is critical for linking overnutrition to the control of neural leptin actions.

## Introduction

Obesity is recognized as a major health concern in children and adults worldwide and is associated with serious comorbidities such as type 2 diabetes, hypertension, cardiovascular disease, certain cancers, and reduced life expectancy. Losing weight or preventing weight gain has a beneficial impact on several metabolic risk factors.

The hypothalamus, located in the central nervous system, is an important site that controls energy balance and maintains normal weight and feeding behavior by sensing hormones that regulate nutrients and appetite. Leptin is the key hormone that acts on the hypothalamus through the leptin receptor and mediates the maintenance of energy homeostasis and normal body weight. Numerous studies have established the crucial role of leptin in controlling energy balance and body weight in both rodents and humans [1-9]. However, obesogenic conditions, such as high-fat diet (HFD) feeding, cause abnormal responsiveness of the hypothalamus to leptin [4-6].

Dietary intake high in saturated fatty acids, especially palmitic acid, reduces the responsiveness of the hypothalamus to leptin and insulin for the regulation of food intake and body weight [10,11]. In previous studies, we also confirmed that palmitic acid directly inhibits leptin signaling in the hypothalamus, specifically the phosphorylation of STAT3 (p-STAT3), a critical mediator of leptin action, by investigating *ex vivo* hypothalamic brain slice cultures [12-15].

However, in the present study, we found that 2-monopalmitin, the palmitic acid bound to glycerol in the β-position (sn-2), enhances hypothalamic leptin responsiveness in organotypic hypothalamus slices. Notably, 1-monopalmitin failed to show an enhanced effect on leptin signaling. Previous studies have focused mainly on the presence or absence of double bonds, carbon number, and saturation of free fatty acids, but few studies have focused on the position of glycerol bonds. Thus, it is necessary to elucidate the mechanisms mediating the effects of 2-monopalmitin on energy metabolism and obesity.

## Results

### 2-monopalmitin enhances hypothalamic leptin responsiveness in *ex vivo* slices and obese mice

First, we investigated whether 2-monopalmitin affects hypothalamic leptin sensitivity. To examine the potential role of 2-monopalmitin on hypothalamic leptin signaling, we used organotypic hypothalamus slice cultures as an *in vitro* model of cellular leptin signaling. pSTAT3 was observed in control slices stimulated with leptin, as shown in Figure 1A. Whereas leptin-induced pSTAT3 levels were significantly reduced by palmitic acid (100 μM, 24 h), the treatment with 2-monopalmitin (100 μM, 24 h) significantly increased leptin-induced pSTAT3 levels in organotypic hypothalamus slices (Figure. 1A). Notably, we found that 2-monopalmitin (30 μM, 24 h) increased leptin sensitivity, whereas its structural analogs, 1-monopalmitin (30 μM, 24 h) and palmitic acid with glycerol (30 μM, 24 h), failed to increase leptin sensitivity in *ex vivo* slices (Figure 1B). These results suggested that the 2-monopalmitin, but not 1-monopalmitin, treatment increased cellular leptin responsiveness.

**Figure 1.**
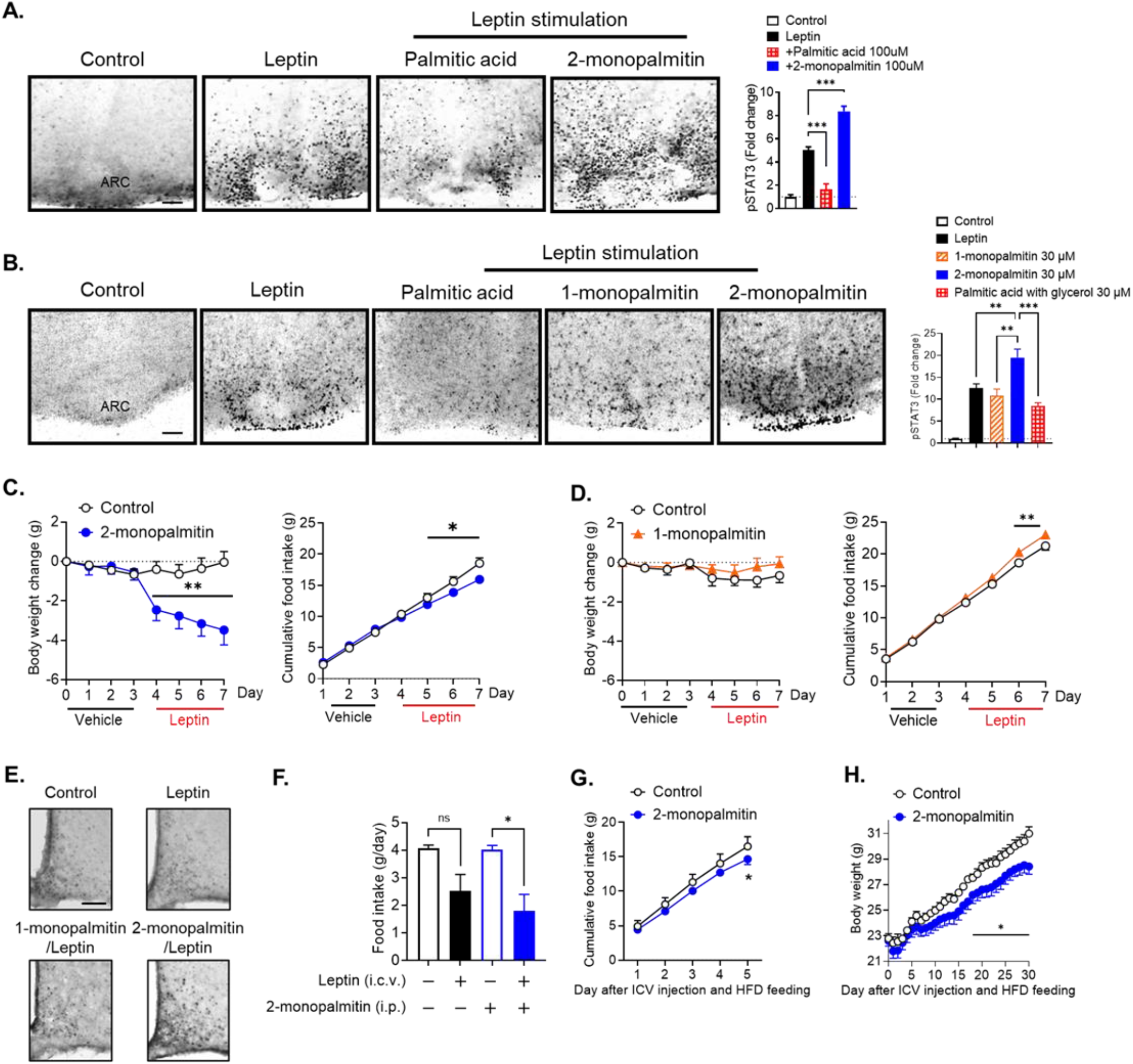
2-monopalmitin enhanced hypothalamic leptin responsiveness and protects from diet-induced obesity. (A) Palmitic acid inhibits central leptin signaling, but 2-monopalmitin enhances central leptin sensitivity under similar conditions. Hypothalamic slice cultures were incubated with 2-monopalmin (100 μM, 24 h) or palmitic acid (100 μM, 24 h) and then stimulated with leptin (30 nM, 60 min). Representative immunohistochemical images of hypothalamic pSTAT3 in slices and quantification of hypothalamic pSTAT3 (n = 6) are shown. Scale bar: 100 μm. (B) 2-monopalmitin, but not 1-monopalmitin or palmitic acid, enhances central leptin responsiveness. Hypothalamic slice cultures were incubated with 2-monopalmitin (30 μM, 24 h), 1-monopalmitin (30 μM, 24 h), or palmitic acid (30 μM, 24 h) and then stimulated with leptin (30 nM, 60 min). Representative immunohistochemical images of hypothalamic pSTAT3 in slices and quantification of hypothalamic pSTAT3 (n = 4) are shown. Scale bar: 100 μm. (C and D) HFD-fed obese mice (n = 5, 4 months of HFD feeding) were i.c.v. administered 2-monopalmitin (2 µg/day), 1-monopalmitin (2 µg/day), or vehicle. Leptin (0.5 μg/day) or vehicle was i.c.v. administered. Body weight and food intake were measured. (E) Mice (n = 3) were i.c.v. administered 2-monopalmitin, 1-monopalmitin, or vehicle followed by leptin (0.5 μg) 3 h later. p-STAT3 immunohistochemistry and quantification. Scale bar: 100 μm. (F) Mice received i.p. injections of 2-monopalmitin (100 µg) twice a day (10 a.m. and 5 p.m.) and then i.c.v. injections of leptin (0.5 μg, 5 p.m.) after the last 2-monopalmitin injection. Food intake was measured 24 h after leptin injection (n = 5–7). (G and H) 2-monopalmitin was centrally infused (2 μg, once a day) into HFD-fed mice. The food intake (G) and body weight change (H) are shown. Each data point represents the mean ± SEM. *p < 0.05, **p < 0.01, and ***p < 0.001 for two-way ANOVA followed by Sidak’s multiple comparisons tests in (C, D, G, and H) and one-way ANOVA followed by Tukey’s multiple comparisons tests in (A, B, and F).

Our results imply that 2-monopalmitin can improve leptin sensitivity in dietary obesity. Therefore, we investigated whether centrally administered 2-monopalmitin increased leptin responsiveness in diet-induced obese mice. Mice were fed an HFD for 3 months, and these age- and weight-matched groups were used to evaluate the anorexic response following the central administration of leptin. 2-monopalmitin or vehicle was centrally administered for 3 days before the central injection of leptin. Markedly, the central injection of leptin resulted in significantly reduced body weight and suppressed food intake in 2-monopalmitin-treated mice, whereas leptin-induced body weight loss and feeding suppression were abolished in vehicle-treated mice (Figure 1C). In contrast to 2-monopalmitin, under these conditions, 1-monopalmitin-treated mice failed to show an anorectic response to exogenous leptin (Figure 1D). Moreover, the central injection of 2-monopalmitin was found to enhance leptin-dependent hypothalamic STAT3 phosphorylation in obese mice, whereas this effect was not observed with 1-monopalmitin (Figure 1D). Significantly, peripheral administration of 2-monopalmitin also enhanced anorectic responses to exogenously administered leptin (Figure 1D). Furthermore, cellular leptin sensitivity, as demonstrated by leptin-induced phosphorylation of STAT3, was significantly enhanced in 2-monopalmitin-treated mice but absent in controls and 1-monopalmitin-treated mice under HFD conditions (Figure 1E). The peripheral injection of 2-monopalmitin enhanced the feeding suppression effect of centrally administered leptin under HFD conditions (Figure 1F).

Next, we investigated whether 2-monopalmitin has an anti-obesity effect when centrally administered alone to HFD-induced hyperleptinemic and leptin-resistant obese mice. Central daily infusion of 2-monopalmitin significantly reduced the food intake (Figure 1G) and body weight gain (Figure 1H) of HFD-fed mice compared with vehicle-treated control mice. Thus, chronic administration of 2-monopalmitin alone is sufficient to decrease body weight under obesogenic conditions (vehicle versus 2-monopalmitin, p < 0.05). Collectively, these findings demonstrate that 2-monopalmitin reverses decreased leptin responsiveness and reduced body weight in mice under hypercaloric feeding.

### 2-monopalmitin affects energy balance

Because the central treatment of 2-monopalmitin resulted in an anti-obesity effect, we further investigated the role of 2-monopalmitin on energy expenditure by directly assessing energy balance in open-circuit indirect calorimetry cages under obesogenic conditions. We observed energy expenditure (oxygen consumption, carbon dioxide production, or heat production) and showed that the parameters pertaining to energy expenditure were slightly increased in 2-monopalmitin-treated mice (Figure 2A–2C). Moreover, 2-monopalmitin-treated mice showed a lower respiratory quotient than vehicle-treated controls, indicating the preferential use of fat as an energy source (Figure 2D). In contrast, no difference in energy expenditure and respiratory quotient was observed between 1-monopalmitin and control mice (Figure 2E–2H). To further investigate the effect of 2-monopalmitin on energy expenditure, we centrally administered 2-monopalmitin into leptin-deficient *ob/ob* mice, another mouse model of obesity. In *ob/ob* mice, 2-monopalmitin did not induce an improvement in energy expenditure (Figure 2I–2L), suggesting that 2-monopalmitin in the brain acts through leptin signaling. Thus, in addition to its role in leptin sensitivity, the enhanced energy expenditure and preferential oxidation of fat as an energy substrate in 2-monopalmitin-treated mice may contribute to the protection against body weight gain under obesogenic conditions.

**Figure 2.**
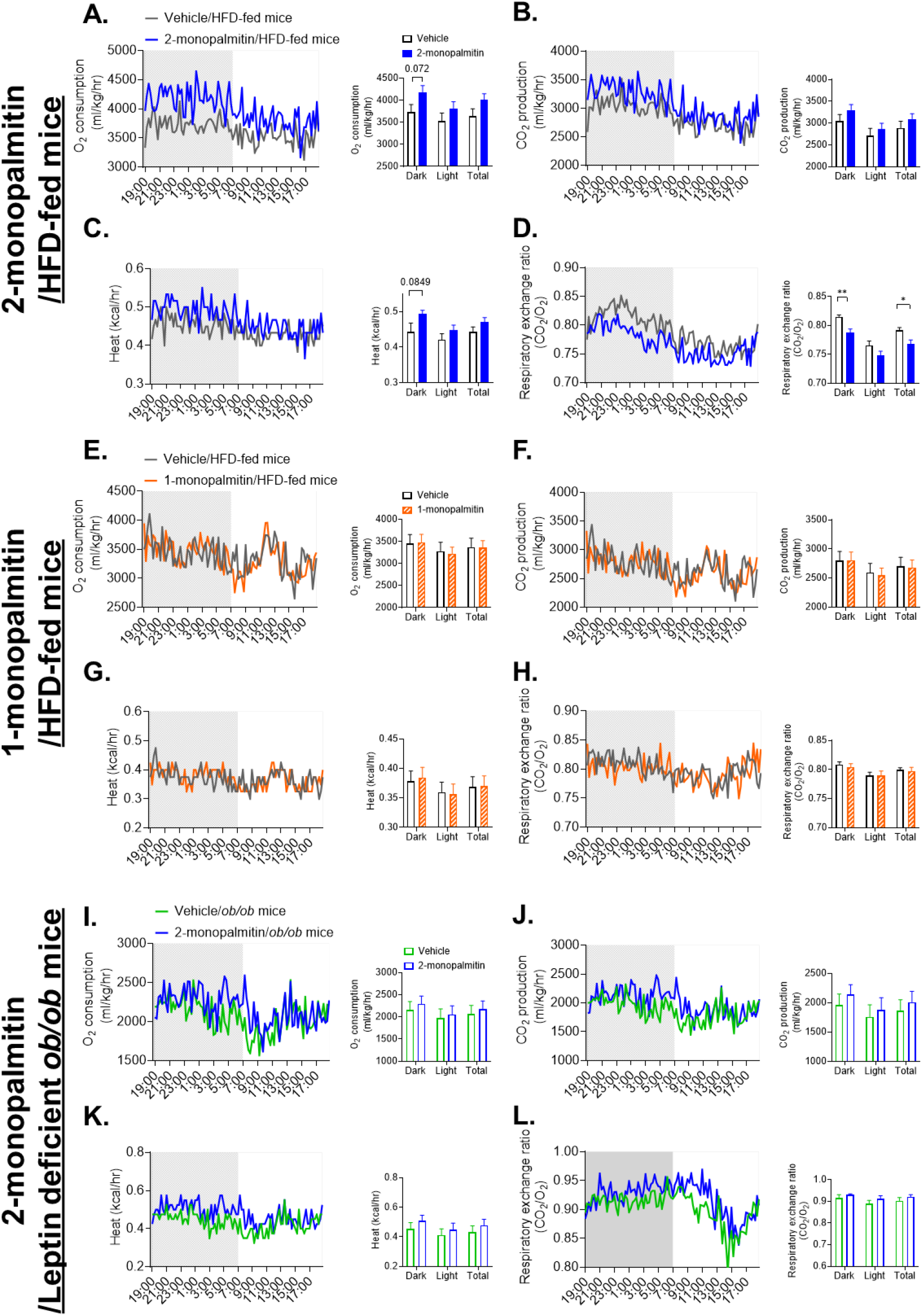
Centrally administered 2-monopalmitin enhances energy balance. (A–D) Metabolic profile of males fed an HFD for 2 weeks with 2-monopalmitin (2 µg/day, i.c.v.) treatment (n = 4/group); O_2_ consumption (A), CO_2_ production (B), heat production (C), and respiratory exchange ratio (D) during 24-h dark or light cycles. The HFD was initiated at 8 weeks of age and mice at 10 weeks of age had comparable body weight (Vehicle: 25.60 ± 0.91 g vs. 2-monopalmitin: 24.61 ± 0.49 g, p > 0.05, t-tests) in the CLAMS study. (E–H) Metabolic profile of males fed an HFD for 4 weeks with 1-monopalmitin (2 µg/day, i.c.v.) treatment (n = 4/group); O_2_ consumption (E), CO_2_ production (F), heat production (G), and respiratory exchange ratio (H) during 24-h dark or light cycles. The HFD was initiated at 8 weeks of age and mice at 10 weeks of age had comparable body weight (Vehicle: 22.77 ± 0.32 g vs. 1-monopalmitin: 23.15 ± 0.58g, p > 0.05, t-tests) in the CLAMS study. (I–L) Metabolic profile of leptin-deficient *ob/ob* obese males with 2-monopalmitin (2 µg/day, i.c.v.) treatment for 2 weeks (n = 4/group); O_2_ consumption (I), CO_2_ production (J), heat production (K), and respiratory exchange ratio (L) during 24-h dark or light cycles. Note that mice at 10 weeks of age had comparable body weight (Vehicle: 42.55 ± 1.24 g vs. 2-monopalmitin: 44.53 ± 0.56 g, p > 0.05, t-tests) in the CLAMS study. Note that there were no differences in ambulatory activity between each experimental group. Each data point represents the mean ± SEM. *p < 0.05 and **p < 0.01 for t-tests in (D).

### Improved glucose balance and peripheral insulin sensitivity in central 2-monopalmitin-treated mice

Consistent with the leaner body weight phenotype, central administration of 2-monopalmitin induced significantly lower blood glucose levels than those observed in vehicle-treated control mice under HFD feeding (Figure 3A). Furthermore, centrally administrated 1-monopalmitin did not affect blood glucose in both the fed and fasted states (Figure 3D). These results suggested that 2-monopalmitin in the brain, but not 1-monopalmitin, increased peripheral insulin sensitivity. Under HFD feeding, 2-monopalmitin-treated mice showed enhanced glucose tolerance (Figure 3B) and improved insulin sensitivity (Figure 3C), whereas 1-monopalmitin-treated mice failed to show an improved glucose balance (Figure 3E and 3F). In further support of the improved insulin sensitivity in 2-monopalmitin-treated mice, insulin signaling was significantly enhanced in the liver and muscle after central administration of 2-monopalmitin, as assessed by western blot analyses using phospho-specific antibodies to Akt (Figure 3G), the central mediators of insulin signaling [16]. Collectively, these findings suggest that in addition to its role in body weight regulation, 2-monopalmitin regulates glucose balance and peripheral insulin sensitivity under obesogenic conditions.

**Figure 3.**
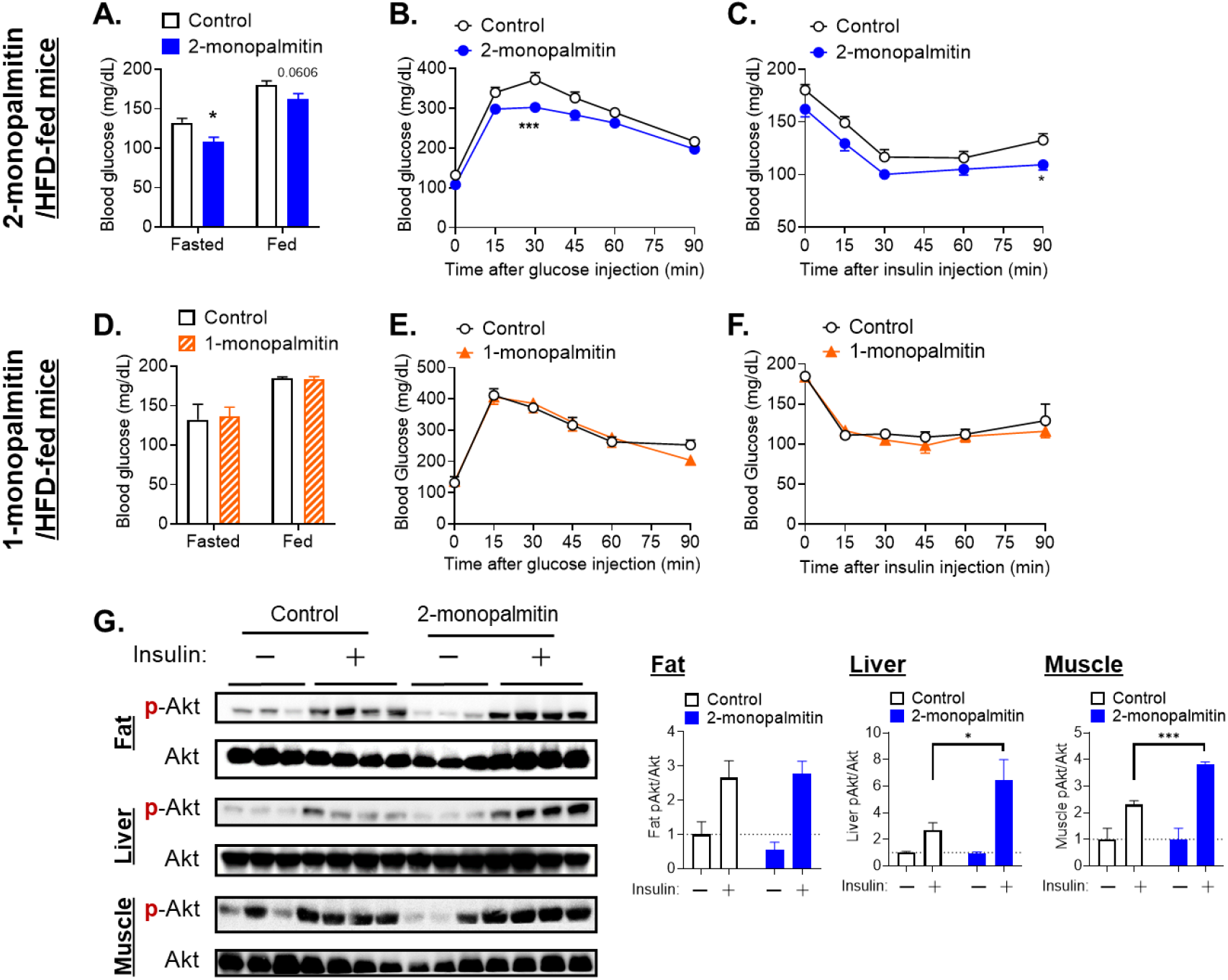
Centrally administered 2-monopalmitin enhances glucose balance in HFD-fed mice. (A–C) Glucose homeostasis parameters of 2-monopalmitin-treated (2 µg/mouse, i.c.v., once a day for 1 week) or control mice fed an HFD for 4 weeks (n = 8–10/group). The glucose (A), glucose tolerance test (GTT) (B), and insulin tolerance test (ITT) (C) are shown. (D–F) Glucose homeostasis parameters of 1-monopalmitin-treated (2 µg/mouse, i.c.v., once a day for 1 week) or control mice fed an HFD for 4 weeks (n = 4/group). The glucose (D), glucose tolerance test (GTT) (E), and insulin tolerance test (ITT) (F) are shown. (I) Cellular insulin sensitivity (n = 3–4/group). The western blot (left) and quantification (right) of Akt (Thr308) phosphorylation in the fat, liver, and muscle 10 min after a bolus injection of insulin (1 U/kg, i.p.) or saline into 2-monopalmitin-treated (2 µg/mouse, i.c.v., once a day for 1 week) or control mice fed an HFD for 4 weeks. Each data point represents the mean ± SEM. *p < 0.05 and ***p < 0.001 for t-tests in (A), two-way ANOVA followed by Sidak’s multiple comparisons tests in (B and C), and one-way ANOVA followed by Tukey’s multiple comparisons tests in (G).

### 2-monopalmitin inhibits hypothalamic inflammation

Because HFD feeding causes inflammation and is involved in decreased leptin responsiveness in the hypothalamus, we next investigated whether 2-monopalmitin regulates cellular processes mediating inflammation to enhance leptin sensitivity. To test this, we used a microglial cell line, namely SIM-A9 cells, and found that treatment with 2-monopalmitin protected them from LPS-mediated inflammation (Figure 4A). In addition, treatment with 2-monopalmitin decreased *Socs-3, IL1β*, and *TNFα* mRNA expression more than treatment with 1-monopalmitin in the hypothalamus of HFD-induced obese mice (Figure 4B). Furthermore, using *ex vivo* slices, we demonstrated that treatment with 2-monopalmitin significantly protected against LPS-mediated decreased leptin responsiveness (Figure 4C). These data indicate that 2-monopalmitin inhibits inflammation and enhances hypothalamic leptin responsiveness in the brain under obesogenic conditions.

**Figure 4.**
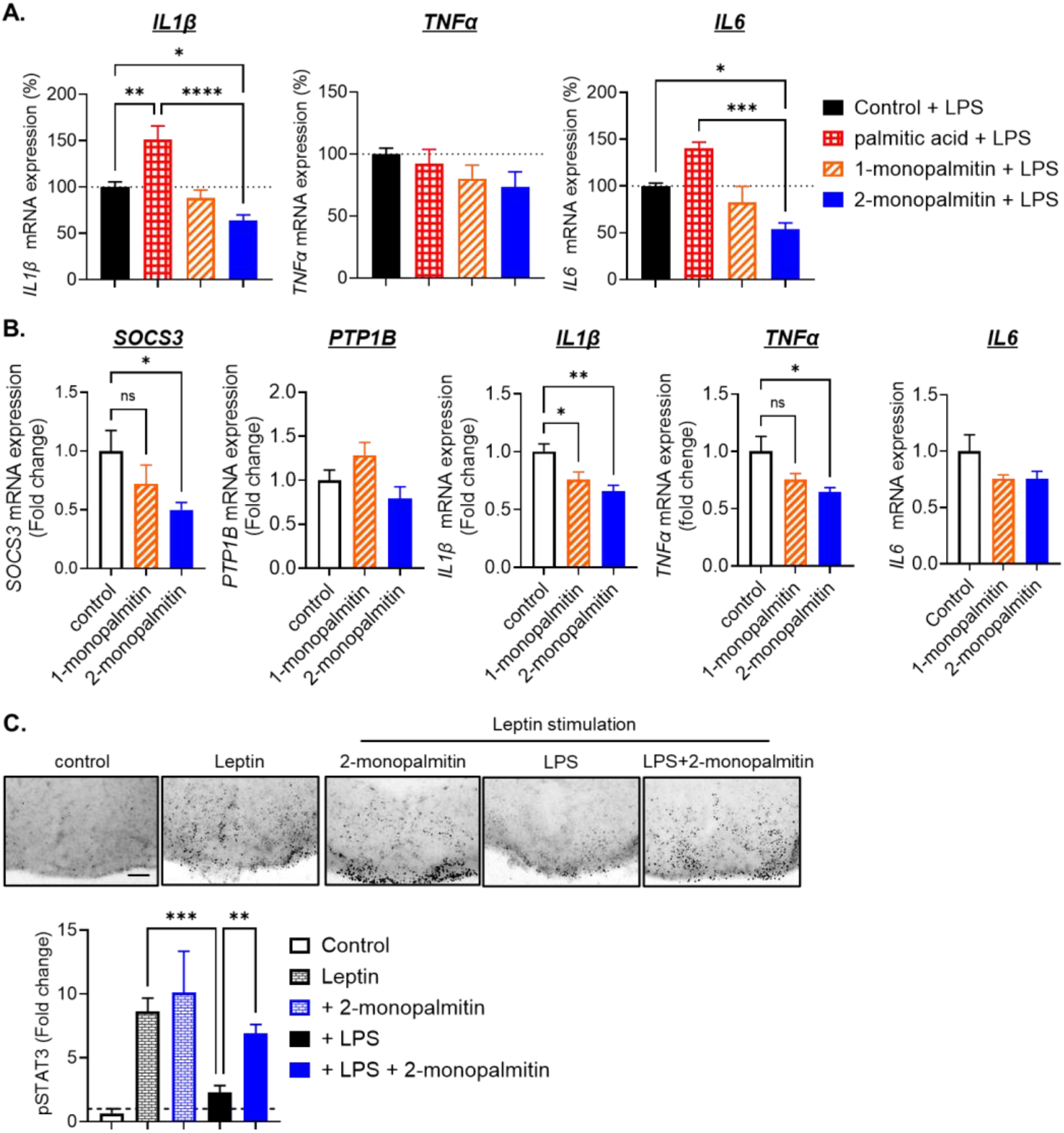
2-monopalmitin decreases hypothalamic inflammation. (A) Relative mRNA expression of *IL-1β, TNFα*, and *IL-6* in SIM-A9 cells. The cells were incubated with vehicle, palmitic acid (100 μM), 1-monopalmitin (100 μM) or 2-monopalmitin (100 μM) for 18h and then stimulated with LPS (0.1 µg/mL, 4 h). Data are from two different experiments (n = 4–7). (B) Hypothalamic expression of genes involved in leptin resistance in 2-monopalmitin-treated (2 µg/mouse, i.c.v., once a day for 10 days) or control mice fed an HFD for 4 weeks. Then, 2 h after the last bolus injection, the hypothalami were collected and the indicated mRNAs were examined. (C) Representative immunohistochemical images of hypothalamic pSTAT3 in hypothalamic slices and quantification of hypothalamic pSTAT3 (n = 4). The slices were incubated with LPS (100 ng/mL) in the presence or absence of 2-monopalmitin (30 μM) for 20 h and then stimulated with leptin (60 nM, 60 min). Scale bar: 100 µm. Each data point represents the mean ± SEM. *p < 0.05, **p < 0.01, and ***p < 0.001 for one-way ANOVA followed by Tukey’s multiple comparisons tests in (A, B, and C).

## Discussion

Herein, we showed that 2-monopalmitin, but not 1-monopalmitin, enhances hypothalamic leptin responsiveness. Although it is known that free saturated long-chain fatty acids promote obesity by altering hypothalamic energy homeostatic regulation and inducing hypothalamic inflammation, insulin, and leptin resistance, the majority of fatty acid studies have only considered the biological regulatory effects of free fatty acids based on carbon number and the presence of double bonds. In general, palmitic acid is known to impair the ability of leptin and insulin to regulate food intake and body weight, regardless of obesity [10]. Furthermore, previous studies have shown that an elevated palmitic acid concentration in the brain promotes obesity and obesity-related metabolic disorders by impairing hypothalamic leptin signaling [17,18]. However, few studies have examined the physiological function of palmitic acid as a monoacylglycerol on energy metabolism. In the present study, we provide evidence showing that, unlike 1-monopalmitate, 2-monopalmitate significantly enhances leptin-dependent hypothalamic STAT3 phosphorylation in *ex vivo* brain slices. In addition, centrally administered 2-monopalmiate markedly restored the leptin-induced suppression of food intake and reduction of body weight in diet-induced obese mice. Consistently, 2-monopalmitate administration, not 1-monopalmitate, suppressed hypothalamic expression of SOCS-3 [19-21] and inflammatory cytokines [22-30], known to mediate HFD-induced leptin resistance. These results indicated that different locations of palmitic acid binding in the glycerol backbone affect leptin sensitivity in the hypothalamus.

In addition to its role in leptin sensitivity, the energy expenditure of 2-monopalmitin-treated mice was significantly enhanced compared with controls. 2-monopalmitin-treated mice have a decreased respiratory quotient, suggesting a preferential use of fat as an energy source. Additionally, these results observed with 2-monopalmitin administration were suppressed in *ob/ob* mice, indicating that the increased energy expenditure and decreased respiratory quotient induced by 2-monopalmitin are manifested through leptin’s action. Thus, 2-monopalmitin infusion alone causes protection of HFD-induced body weight gain, likely by enhancing leptin insensitivity. These results demonstrate the potential of 2-monopalmitin as a leptin sensitizer and highlight the importance of the binding position of fatty acids in the glycerol backbone, which was previously unrecognized in anti-obesity research.

Intracerebroventricular administration of 2-monopalmitin improved glucose tolerance, increased systemic insulin sensitivity, and increased cellular insulin signaling in skeletal muscle and liver under HFD feeding. The molecular mechanisms mediating these effects are unknown but may involve a decrease in hypothalamic SOCS-3 because selective deletion of SOCS-3 in the central nervous system improves glucose tolerance and insulin tolerance and increases peripheral insulin sensitivity [31,32]. However, additional mechanisms are likely involved, and further studies are necessary to determine how 2-monopalmitin affects systemic glucose balance.

Human milk fat is known to be high in 2-monopalmitin; palmitic acid accounts for approximately 20%–25% of the fatty acids in human milk. Notably, the majority of palmitic acid in human breast milk is esterified to the sn-2 position of triacylglycerol [33-38], and absorption of palmitic acid increases when it is bound to the sn-2 position of the glycerol skeleton [39,40]. For infants, the increased absorption of 2-monopalmitin suggests that it is an important source of energy metabolism. Considering other natural fats, both lard and beef tallow contain approximately 24% palmitic acid, but more than 70.5 mol% of the palmitic acid in lard is located in the sn-2 position, whereas only ∼15 mol% of the palmitic acid in beef tallow is in the sn-2 position [41-43]. We have recently shown that body weight gain during overnutrition is reduced in lard-fed mice compared with beef tallow-fed mice and that hypothalamic leptin sensitivity is maintained higher in lard-fed mice than in beef tallow-fed mice (*submitted*). Although the mechanism by which 2-monopalmitin is delivered to the brain remains unclear, we have shown that central administration of 2-monopalmitin increases energy expenditure, decreases respiratory quotient, and improves leptin and insulin sensitivity. Most importantly, peripheral injection of 2-monopalmitin enhanced the anorectic effect of centrally administered leptin. Taken together, these findings suggest that 2-monopalmitin may plays a role in maintaining normal body weight in infants and adults.

Our *ex vivo* studies revealed a previously unidentified link between 2-monopalmitin and inflammation in the brain, the relevance of which was substantiated in microglial cells *in vitro* and *in vivo*. Although these findings predict that the inflammation induced by HFD subsequently leads to leptin resistance, 2-monopalmitin can prevent the increased inflammatory response induced by a high-fat diet. Similarly, centrally administered 2-monopalmitin significantly reduced SOCS-3, which is upregulated in the hypothalamus by HFD-induced obesity and is known to be a direct inhibitor of leptin signaling. These results may help clarify its effects on leptin sensitivity and energy balance. Moreover, these findings strongly support the view that the effects of 2-monopalmitin alone are robust enough to restore leptin sensitivity under obesogenic conditions.

In the present study, we considered only the context of HFD-induced hypothalamic metabolic abnormalities, such as leptin and insulin resistance. However, we found that 2-monopalmitin decreases the expression of inflammatory cytokines and SOCS3 in the hypothalamus under HFD conditions. Indeed, hypothalamic inflammation is induced by and causally associated with other pathophysiological conditions. In particular, hypothalamic inflammation occurs with aging and is indicated by the increased gene expression of inflammatory cytokines and inflammation-related signals [44,45]. Given that 2-monopalmitin protects against the induction of inflammatory cytokines in the hypothalamus, 2-monopalmitin may be effective in ameliorating age-related conditions. Whether 2-monopalmitin can be used to improve age-related dysfunction (e.g., emotional, sleep, and cognitive function) is unclear and should be addressed in future research.

In summary, our results suggest that 2-monopalmitin, but not 1-monopalmitin, plays an important role in the protection against HFD-induced decreased leptin responsiveness. The findings also revealed the anti-obesity and anti-diabetic effects of 2-monopalmitin under overnutrition conditions. Thus, we propose that 2-monopalmitin may be valuable in therapeutic interventions to alleviate obesogenic conditions.

## Materials and Methods

### Animals and diet

Male C57BL/6 mice (3 or 7 weeks old) and *ob/ob* mice were purchased from Japan SLC (Shizuoka, Japan). Mice were housed under constant temperature (22–24°C) and humidity (50%–60%), using 12-h light and dark cycles (lights on 7 a.m.–7 p.m.). Mice were fed an HFD (60 kcal% fat, Research Diet, D12492) with *ad libitum* access to water. All experimental procedures were conducted following the ARRIVE guidelines. The care of all animals and the procedures were approved by the Meiji University Animal Committee (MUIACUC2022-05).

### Organotypic hypothalamus slice culture

The organotypic slice culture was performed as previously described [7,8,11]. Hypothalamic slices were obtained using a previously reported method with slight modifications [7,8,11]. C57BL/6 pups at 8–11 days old were decapitated, and the brains were quickly removed. Hypothalamic tissues were sectioned at a depth of 250 μm using a vibratome (Leica VT1200S) in chilled Gey’s balanced salt solution enriched with glucose (0.5%) and KCl (30 mM). Coronal slices containing the arcuate nucleus (ARC) were then placed on Millicell-CM filters (Millipore, pore size of 0.4 μm, diameter of 30 mm) and maintained at an air medium interface in minimum essential media supplemented with heat-inactivated horse serum (25%), glucose (32 mM), and GlutaMAX (2 mM). Cultures were typically maintained for 8– 10 days in standard medium, which was replaced 2 or 3 times a week. Healthy slices typically showed a slight reduction in the thickness of the hypothalamus after 10 days of incubation, whereas marked decreases were observed in unhealthy slices, which mostly lost the hypothalamic structure. After 10 days, the selected healthy slices were randomly divided into experimental and control groups.

### Immunohistochemistry

Immunohistochemistry was performed as previously described [7]. Under deep anesthesia, male mice were intracardially perfused with saline and 4% paraformaldehyde. The brains were removed, post-fixed in 4% paraformaldehyde, infiltrated with 20% sucrose, and cut into 30 μm-thick slices. Sections were rinsed 6 times for 5 min each in PBS and then placed in 0.3% hydrogen peroxide and 0.25% Triton X-100 in PBS (PBT) for 30 min. Next, sections were incubated for 48–72 h with phosphorylated STAT3 (pSTAT3) antibodies (1:3000, Cell Signaling Technology, 9131) in 3% normal donkey serum with PBT and 0.02% sodium azide. Sections were reacted with a biotinylated secondary antibody against rabbit IgG (1:1000, Jackson ImmunoResearch Labs, West Grove, PA) followed by the avidin-biotin-peroxidase complex (ABC) kit (1:1000, Vectastain Elite ABC kit; Vector Labs, Burlingame, CA). Immunoreactivities were visualized by incubation with 3,3-diaminobenzidine (DAB, Sigma, St. Louis, MO). After dehydration through a graded series of ethanol, slides were immersed in xylene and coverslipped. Images were captured using a brightfield Nikon ECLIPSE Ts2 microscope. Organotypic hypothalamic slices were cut from the membrane, rinsed 3 times for 10 min each in PBS at pH 7.4, and then placed for 20 min in 1% hydrogen peroxide and 1% sodium hydroxide in PBS to quench endogenous peroxidase activity. Following a series of washes with PBS, slices were incubated at 4°C for 48–72 h in pSTAT3 antibodies (1:3000) in 3% normal donkey serum (Jackson ImmunoResearch Labs) with PBT and 0.02% sodium azide. After washing in PBS, slices were incubated with a biotinylated donkey anti-rabbit antibody (1:1000) in 3% donkey serum in PBT at room temperature for 1 h. Tissues were rinsed in PBS and incubated in ABC (1:500) for 1 h. Slices were washed in PBS and reacted with DAB, and then rinsed in PBS and mounted on slides using Vectashield (Vector Labs). To measure the pSTAT3 signal in the ARC, the uneven background was eliminated using Adobe Photoshop, and the intensity of DAB staining was measured using NIH ImageJ.

### Energy expenditure and food intake measurements

Mice were first acclimatized to the metabolic cages and housed individually for 2–3 days before measurements were taken. Metabolic parameters, including O_2_ consumption, CO_2_ production, respiratory exchange ratio, heat production, and ambulatory activity were determined using a comprehensive lab animal monitoring system (CLAMS) (Columbus Instruments, Ohio, USA), as previously described [7]. Ambulatory activity was measured using an ACTIMO system (Shinfactory, Fukuoka, Japan).

### Cannula implantation and leptin sensitivity test

The cannula implantation was performed as previously described [7-9,11]. Mice were anesthetized with isoflurane and positioned in a stereotaxic frame. A 26-gauge single stainless-steel guide cannula (C315GS-5-SPC, Plastics One, Roanoke, VA, USA) was implanted into the lateral ventricles (−0.45 mm from the bregma, ± 0.9 mm lateral, and −2.5 mm from the skull). The cannula was fixed to the skull using screws and dental cement. The mice were housed in single cages and allowed to recover from the operation for 1 week. At the end of the experiment, the placement of the guide cannula was histologically verified. For the leptin sensitivity test, after 4 months of HFD feeding, an ICV cannula was implanted, and mice were allowed 1 week to recover. On experimental days, as previously described [7-9,11], 2-monopalmitin (2 µg/mouse, Cayman Chemical 17882), 1-monopalmitin (2 µg/mouse, TCI G0083), or DMSO was administered at 5 p.m. for 3 days, and mice were then treated with leptin (0.5 μg/mouse, PeproTech) or saline once a day at 5 p.m. for 3 consecutive days. Food intake and body weight were measured daily. For peripheral injection, 2-monopalmitin was dissolved in 0.45% (2-hydroxypropyl)-beta-cyclodextrin solution.

### Glucose and insulin tolerance tests

Blood glucose levels were determined in freshly withdrawn blood from the tail vein using a OneTouch Ultra blood glucose meter. Glucose tolerance tests were performed on overnight-fasted mice. D-glucose (1.5 g/kg) was injected intraperitoneally, and blood glucose was measured at the indicated periods from the tail vein. Insulin tolerance tests were performed on 3 h-fasted mice. Insulin (1.0 U/kg) was injected intraperitoneally, and blood glucose was measured at the indicated periods.

### Total protein extraction and western blot analysis

Western blot analysis was performed as previously described, with slight modifications [7-9,11]. Briefly, proteins were extracted by homogenizing samples in lysis buffer (25 mM Tris-HCl at pH 7.4, 150 mM NaCl, 1% NP-40, 1 mM EDTA, and 5% glycerol) (87787 and 87788 Pierce IP Lysis Buffer, Thermo Fisher Scientific) with protease and phosphatase inhibitor cocktails (1:100, 78442, Thermo Fisher Scientific). The hypothalamus was collected as follows: after brief anesthetization with isoflurane, mice were decapitated, and the whole brain was removed. The hypothalamus was prepared using a brain matrix (1 mm thick), frozen immediately in liquid nitrogen, and stored at –80°C. Equal amounts of the samples were separated by SDS-PAGE and transferred to a nitrocellulose membrane by electroblotting. The following primary antibodies were used for western blot assays: a phosphorylated Akt antibody (1:1,000, Cell Signaling Technology, 4060) and an Akt antibody (1:1,000, Cell Signaling Technology, 2920). After incubation with primary antibodies for 72 h at 4°C, the membranes were incubated with gentle agitation for 1 h at room temperature using the following secondary antibodies conjugated to a chemiluminescence entity: anti-rabbit IgG and HRP-linked antibodies (1:5000, Cell Signaling Technology, 7074). WSE-6100 Lumino-GraphI (ATTO Corporation, Tokyo, Japan) was used to measure the chemiluminescence intensity.

### Total RNA extraction and quantitative real-time PCR

Quantitative real-time PCR analysis was performed using a previously reported method with slight modifications [7-9,11]. Hypothalamic samples were collected from C57BL/6 mice fed with lard or beef tallow for 5 months, and total RNA was extracted using the RNeasy Mini Kit (QIA-GEN, Hilden, Germany). cDNA was generated using the Takara Prime Script® RT Master Mix (Takara, Osaka, Japan). Regarding quantitative real-time PCR, we amplified cDNA using the CFX Connect Real-Time PCR Detection System (Bio-Rad Laboratories, Inc. California, USA) with the THUNDERBIRD® qPCR Mix (Toyobo Co., Osaka, Japan), and each primer was set specific for mouse cyclophilin, SOCS-3, protein tyrosine phosphatase 1B (PTP1B), IL-6, IL-1β, and tumor necrosis factor-α (TNFα), performed according to the manufacturer’s instructions, as previously described. Reactions were cycled 40 times for denaturation at 95°C for 15 s and annealing and elongation at 60°C for 60 s. Normalized mRNA levels were expressed in arbitrary units, obtained by dividing the averaged efficiency-corrected values for sample mRNA expression by that for β-actin RNA expression for each sample. The resulting values were expressed as fold change above-average control levels. The following primer sequences were used: SOCS-3 (F-CACCTGGACTCCTATGAGAAAGTG and R-GAGCATCATACTGATCCAGGAACT), PTP1B (F-GGAACAGGTACCGAGATGTCA and R-AGTCATTATCTTCCTGATGCAATT), IL-6 (F-TAGTCCTTCCTACCCCAATTTCC and R-TTGGTCCTTAGCCACTCCTTC), IL-1β (F-GCAACTGTTCCTGAACTCAACT and R-ATCTTTTGGGGTCCGTCAACT), TNFα (F-CCCTCACACTCAGATCATCTTCT and R-GCTACGACGTGGGCTACAG), or β-actin (F-CTGCGCAAGTTAGGTTTTGTCA and R-TGCTTCTAGGCGGACTGTTACTG).

### SIM-A9 cells culture

SIM-A9 cells were purchased from the American Type Culture Collection (ATCC CRL-3265TM, Manassas, VA, USA). SIM-A9 cells were cultured in DMEM/F12 (Gibco, Carlsbad, CA, USA) (pH 7.4) supplemented with 10% (v/v) fetal bovine serum (Hyclone Laboratories Inc., Chicago, IL, USA), 5% (v/v) horse serum (Gibco, Carlsbad, CA, USA), and 1% (v/v) penicillin-streptomycin (Gibco, Carlsbad, CA, USA) at 37°C under 5% CO_2_. On day 1, SIM-A9 was seeded at 8×10^4^ cells/ml in a 24-well plate. Then, the cells were incubated for one day. Each sample of 100 µM concentration was added on the third day, and 18 h later, a mixture of LPS (0.1 µg/mL) and each sample was added. After 4h, the cells were collected using Buffer RLT, sonicated, and used for qPCR.

### Statistical analysis

All data are expressed as means ± SEM. Statistical analysis was performed by employing a two-tailed unpaired Student’s t-test or one- or two-way ANOVA followed by post hoc Tukey’s or Sidak’s tests. All statistical analyses were performed using Prism 10 (GraphPad Software, CA). Herein, p < 0.05 was considered significant.

## Study approval

All procedures used to maintain and evaluate the mice followed protocols reviewed and approved by the Animal Research Committee of Meiji University (MUIACUC2022-05) (Kanagawa, Japan). Animal experimentation guidelines were followed.

## Acknowledgments

The authors gratefully acknowledge Dr. Junko Tanaka and Dr. Keisuke Furuichi for helpful discussion during this study. This study was partially supported by the JST FOREST Program, Grant Number JPMJFR225C to K.K.

## Author contributions

K.K. conceived the study. N.T., M.I., and K.K. designed the experiments. N.T., M.I., Y.Y., Y.F., T.S., A.H., and K.K. performed the experiments, analyzed data, and interpreted the results. The majority of the manuscript was written by K.K., with some help from N.T., M.I., Y.Y., and Y.F. All authors approved the final version of the manuscript. The order of the co-first authors was determined by their relative contribution to this study.

## Data availability

The datasets used and analyzed during the current study are available from the corresponding author upon reasonable request.

